# DeepERA: deep learning enables comprehensive identification of drug-target interactions via embedding of heterogeneous data

**DOI:** 10.1101/2023.01.27.525827

**Authors:** Le Li, Shayne D. Wierbowski, Haiyuan Yu

## Abstract

Drug-target interaction prediction is a crucial step in drug development, discovery, and repurposing. Due to the tremendous combinatorial search space of all drug-protein pairs, machine learning algorithms have been utilized to facilitate the identification of novel drug-target interactions. Deep learning, known as a powerful learning system, has recently shown superior performance to traditional machine learning in many biological and biomedical areas. In this paper, we proposed an end-to-end deep learning model, DeepERA, to identify drug-target interactions based on heterogeneous data. This model assembles three independent feature embedding modules (intrinsic embedding, relational embedding, and annotation embedding) which each represent different attributes of the dataset and jointly contribute to the comprehensive predictions. This is the first work that, to our knowledge, applied deep learning models to learn each intrinsic features, relational features, and annotation features and combine them to predict drug-protein interactions. Our results showed that DeepERA outperformed other deep learning approaches proposed recently. The studies of individual embedding modules explained the dominance of DeepERA and confirmed the effects of the “guilt by associations” assumption on the performance of the prediction model. Using our DeepERA framework, we identified 45,603 novel drug-protein interactions for the whole human proteome, including 356 drug-protein interactions for the human proteins targeted by SARS-CoV-2 viral proteins. We also performed computational docking for the selected interactions and conducted a two-way statistical test to “normalize” the docking scores of different proteins/drugs to support our predictions.

## Introduction

Chemical compounds and small-molecule drugs can inhibit the functions of their target proteins by blocking their active sites or inducing conformational changes that prevent recognition of substrates or other protein interactors. The mechanisms of these drug-target interactions underly the potential efficacy or side-effects of drugs making their identification a critical step in drug development and repurposing. With more than 90 million possible drug candidates (small molecules) and ∼20,000 human proteins (Rifaioglu et al., 2019), the identification of drug-target interactions among this large-scale search space through pure chemical or biological assays is a very time- and resource-consuming procedure (Kapetanovic, 2008). Therefore, computational approaches have been applied to prioritize high-confidence candidates and facilitate the development of drugs.

Traditional computational approaches (e.g. virtual screen, reverse virtual screen) utilized molecular docking technique (A. C. Cheng et al., 2007; Oleg Arthur J., 2010), which rely on 3D structures of proteins that may often be unavailable, or if available may not have captured the correct conformational state to accommodate ligand docking. To overcome this problem, many machine learning approaches predicting drug-target interactions based on 1D or 2D chemical and genomic features have been proposed. These methods can be grouped into two categories according to the features they used: the approaches based on intrinsic features and the approaches based on relational features. The former category focuses on the curation of features of each drug and each protein, such as substructure fingerings for drugs and sequence motifs for proteins(Cao et al., 2012; Ezzat et al., 2018; Mousavian & Masoudi-Nejad, 2014; Nagamine & Sakakibara, 2007; Yamanishi et al., 2011). Relational features are commonly represented as similarities between drugs/proteins or networks/graphs in the studies and have been applied in drug-drug (Qian et al., 2019; Zitnik et al., 2018), drug-disease (H. Luo et al., 2016), protein-protein (Kovács et al., 2019), and drug-protein interaction prediction studies (F. Cheng et al., 2012; Liang Yu, 2020; Madhukar et al., 2019; Mei et al., 2013; van Laarhoven et al., 2011; Yamanishi et al., 2008). These machine learning models have achieved decent performance in the past decade and were the most popular choices for predicting drug-target interactions before the application of deep learning techniques in this field.

Deep learning is a subfield of machine learning that relies on multi-layer artificial neural network architectures to gradually abstract features of data from the raw input. With the rapid development of theoretical research about deep learning and the fundamental computer hardware technologies which provide high-performance computational power, deep learning has been widely applied in biological and biomedical research (including drug-target interaction prediction) in this decade. The earliest deep learning models used within this field had not yet been fully utilized for informed feature generation (i.e., feature embeddings). For example, Tian et al. manually extracted the basic substructures of drugs and the domains of proteins as features and concatenated them to form the feature vectors of drug-protein pairs and fed both positive and negative drug-protein pairs into a deep neural network (DNN) model to predict the interactions (Tian et al., 2016). Some other works extended this study by replacing the curated features with different molecular descriptors (Hamanaka et al., 2017; You et al., 2019) and transcriptional response (Xie et al., 2018) or replacing the DNN architecture with a deep-belief network (DBN) (Wen et al., 2017). More recently, more systematical approaches that generate learned embeddings of protein/drug features have been proposed (Lee et al., 2019; Nguyen et al., 2019; Öztürk et al., 2018, 2019; Tsubaki et al., 2019; Yingkai Gao et al., 2018). This new category of deep learning approaches integrated classification and embedding of drug/protein features into an end-to-end model, thus enabling the simultaneous optimization of feature embedding and prediction modules towards the same training objective while eliminating the need for any a priori domain knowledge or feature engineering. Öztürk et al. embedded the SMILES (Simplified molecular-input line-entry system) strings of drug molecules and protein sequence with CNN (convolutional neural networks) and the concatenated these embeddings to predict the interactions with fully connected (FC) layers. This group extended their work by additionally incorporating LMCS (ligand max. common substructure) and protein motifs and domains in a wider embedding framework (Öztürk et al., 2019). Tsubaki et al. replaced the embedding model of the 2D structure of drug compounds with a novel graph-based neural network (GNN) and made use of the attention mechanism to investigate the drug-binding sites of protein targets (Tsubaki et al., 2019). Gao et al. (Yingkai Gao et al., 2018) further changed the embedding of protein sequence by incorporating GO (gene ontology) annotations of proteins into a long short-term memory based recurrent neural network and increased the model ‘s biological interpretability by utilizing a two-way attention mechanism to dig out drug-binding sites of proteins as well as the contributing weights of compound atoms in the interactions.

The effectiveness of intrinsic embedding has been demonstrated in prior deep learning-based work. However canonical machine learning algorithms have also demonstrated the value of relational information that can incorporate and link disparate prediction tasks such as drug-target interaction and drug-drug side effect predictions (Schlichtkrull et al., 2018; Zitnik et al., 2018). Moreover, the external attributes of drugs and proteins annotated manually by experts or through experimental characterization (i.e. protein-associated phenotypes and diseases, drugs’ side-effects, proteins’ domain information) have been widely applied in traditional machine learning based drug-target interaction prediction works (Y. Luo et al., 2017; Wan et al., 2019) and other topics (e.g. drug repositioning (Zeng et al., 2019)). Because deep learning models were initially introduced to learn the representation of proteins and drugs directly from protein sequences and chemical formulas in predicting drug-target interactions, the relational features and external attributes of drugs and proteins have been largely underutilized in deep learning-based drug-target interaction prediction efforts. The fact that many structurally related drugs bind different targets whereas distinct drugs display significant target overlap (Hu & Bajorath, 2012a) (∼14% of drugs interacting with a given target are structurally similar and ∼72% of structurally related drugs have no or <20% target overlap (Hu & Bajorath, 2012b)) indicates that having structure data (intrinsic features) alone is not sufficient to provide thorough predictions. Recently, some deep learning models incorporating relational features have been proposed (Peng et al., 2021; Zhao et al., 2021). However, these works specifically focused on relational embedding and the information of intrinsic structures proteins and drugs and external annotations were not adequately incorporated into the deep learning architecture. With this motivation, we proposed a <u>Deep</u> learning <u>E</u>mbedding architecture incorporating <u>R</u>elational features and <u>A</u>nnotations of drugs and proteins (DeepERA) (Fig. 1), which combined the advantages of deep learning and heterogeneous data sources, to predict drug-target interactions.

**Fig. 1:**
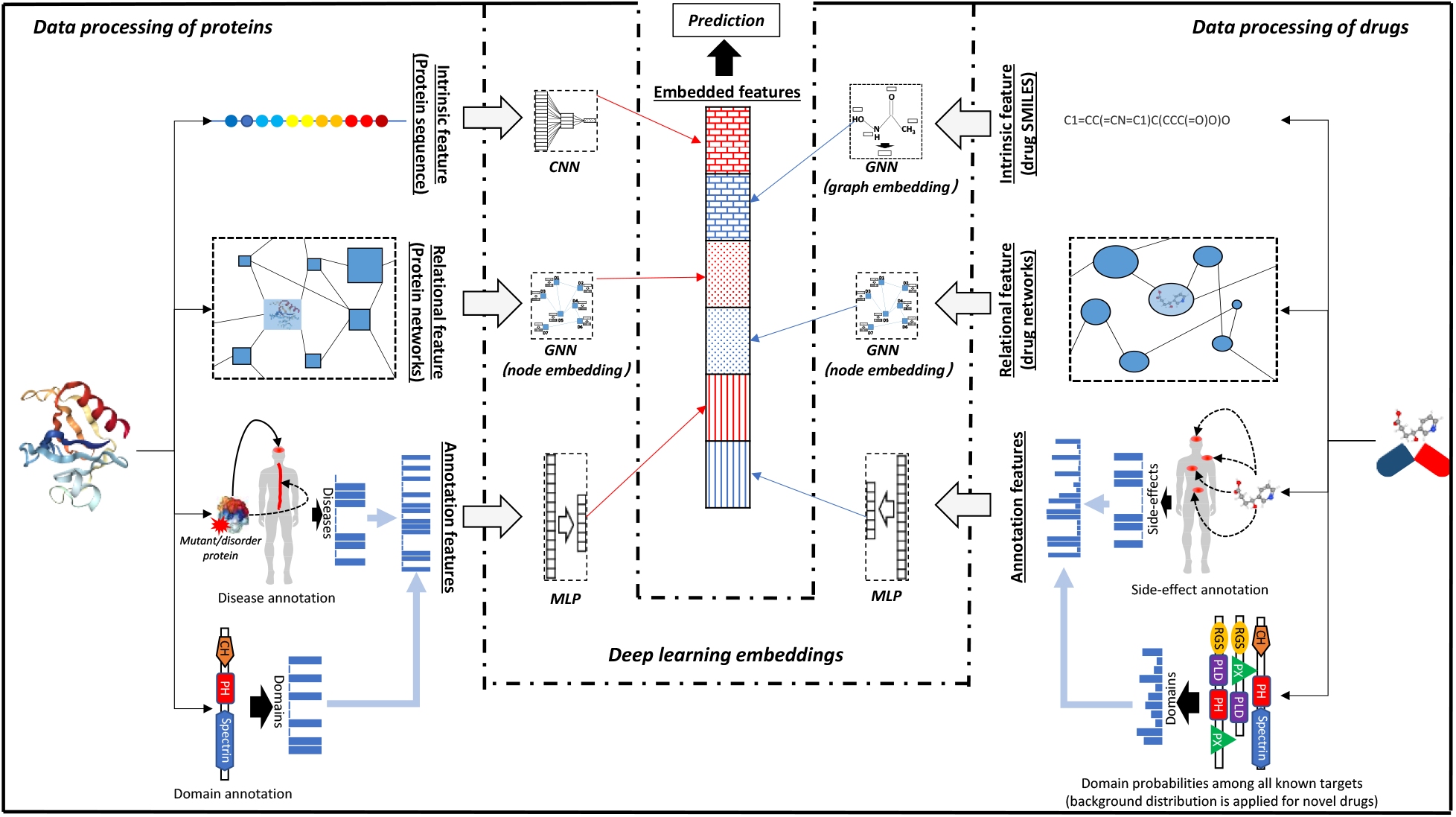
The architecture of DeepERA to predict interaction between drugs and proteins. For each protein and drug, DeepERA processes their intrinsic features (sequence/SMILES strings), relational features (relationship in protein-protein/drug-drug networks), and annotation features (disease/side effects and domain profile) with specific models and the processed features are integrated to make the final prediction.

Considering the natural formats of intrinsic data (protein sequences and drug SMILES strings), relational data (drug networks and protein networks), and external annotations (diseases/side-effects and domains associated with proteins and drugs), we designed three embedding components which learn the features from these three types of data without data transformation procedures to avoid potential information loss. We compared the performance of our approach with the existing deep learning embedding approaches and showed that the proposed approach outperformed the existing deep learning intrinsic embedding approaches. Furthermore, we investigated the contributions of each component in DeepERA in terms of the prediction performance and found that the relational embedding is more capable of uncovering new associations between known drugs targeting unseen proteins than intrinsic embedding and annotation embedding. We also generated a list of potential drugs, based on the predictions of DeepERA, for the human proteins interacting with SARS-CoV-2 viral proteins (Gordon et al., 2020) and provided computational docking with the significance of results determined by statistical tests to prioritize the predictions.

## Result

### DeepERA integrates intrinsic and relational features, and external annotations of both drugs and proteins

The proposed DeepERA architecture (Fig. 1) was fed with a range of protein and drug data types including sequences, SMILES strings, networks, and annotations, and, according to specific properties of each data type, adopted different deep learning modules to learn their representations. The data we used in deep learning embedding can be roughly classified into three categories: intrinsic features, relational features, and external annotated features:

<u>Intrinsic features</u> include protein sequences and drug SMILES strings which represent the most intrinsic attributes of proteins and drugs, and we applied graph neural network (GNN) and convolutional neural network (CNN) to study the embedding of drug SMILES strings and protein sequences, respectively.

<u>Relational features</u> are represented as protein networks and drug networks which denote the interactions or relationships between drugs/proteins, and we learned the relational embedding of the networks with GNN.

<u>Annotation features</u>: disease/side-effect annotations, and Pfam domain annotations of drugs and proteins are curated from other research or experiments which indicate their functions and structure, and they were embedded via multi-layer perceptron (MLP).

All feature vectors embedded from different data sources were concatenated to be the final embedded features of drug-target pairs, which were subsequently fed into a fully connected layer (FC) model to predict if the pairing drugs and proteins interact. In this way, DeepERA learned comprehensive representations of drugs and proteins based on their own properties, their relations with other drugs and proteins, and external annotations to make accurate predictions of drug-protein interactions.

### Integrating multiple types of deep learning-based embeddings based on heterogeneous data improves drug-target interaction prediction

To learn the features from the heterogenous data, DeepERA specified three embedding components based on the formats and properties of the corresponding data: protein sequences and drug SMILES strings are processed by a CNN and a whole-graph GNN, respectively, in the intrinsic embedding component; protein networks and drug networks are processed by single-node GNNs in the relational embedding; external feature vectors are processed by fully connected layers in the annotation embedding. With the studied features from various types of information, DeepERA enabled better prediction of drug-protein interactions.

To investigate the improvement of DeepERA compared with intrinsic-embedding-only deep learning approaches, we compared the performance of DeepERA model with three state-of-the-arts deep learning-based intrinsic embedding methods, i.e., DeepCPI, DeepDTA, DeepDTI. Fig. 2a shows the comparison between DeepERA and these previous approaches for the interactions between unseen drugs and proteins (Type 1), interactions between seen drugs and unseen proteins (Type 2), interactions between unseen drugs and seen proteins (Type 3), and interactions between seen drugs and proteins (Type 4).

**Fig. 2:**
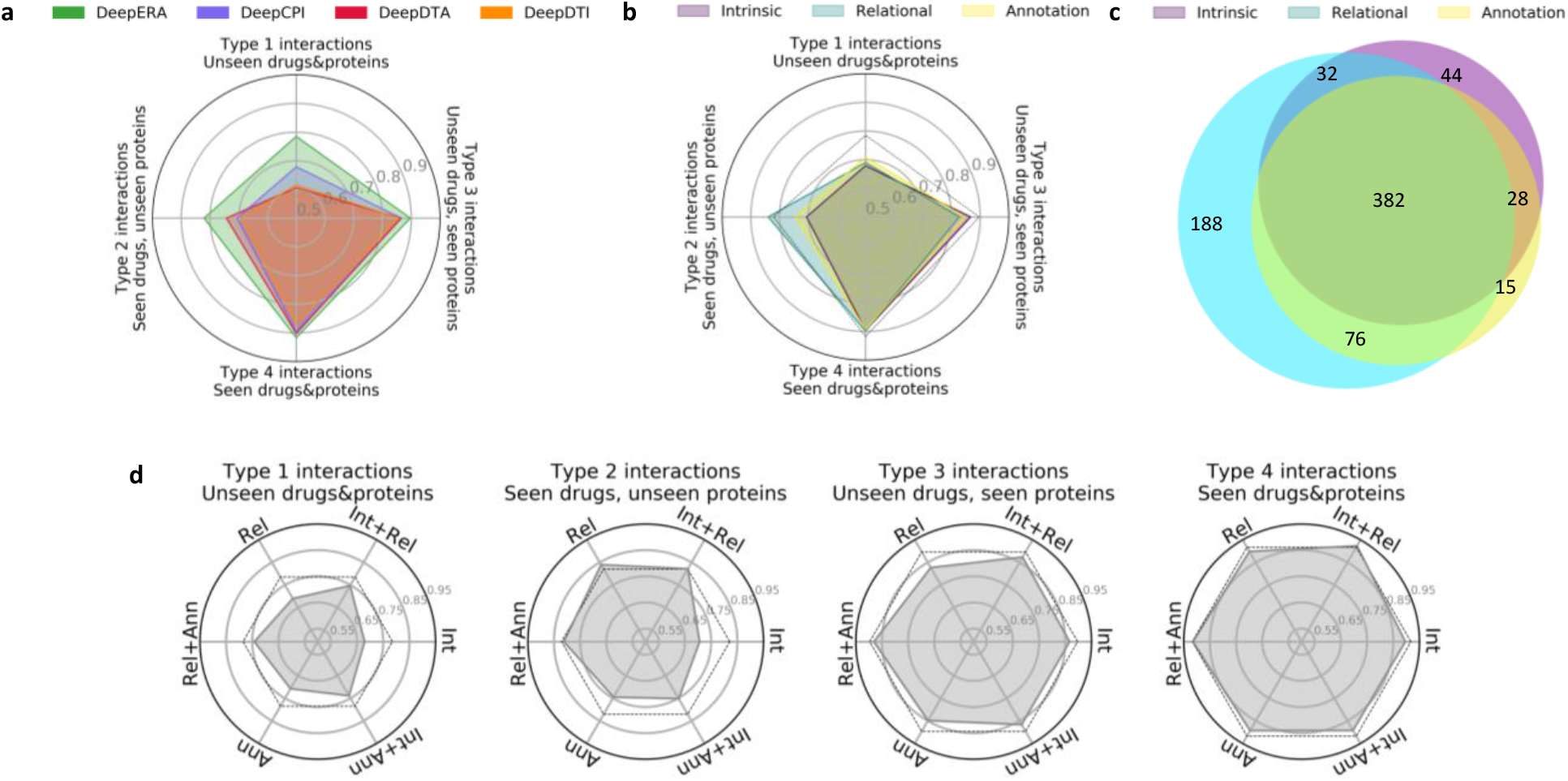
Investigate the performance of DeepERA. **a** Comparison of AUCs between DeepERA and other deep learning approaches. **b** Comparison of AUCs between individual embedding components in DeepERA. Int: intrinsic embedding; Rel: relational embedding; Ann: annotation embedding. **c** Venn diagram of true interactions predicted by different embedding components for the interactions between seen drugs and unseen proteins. **d** AUC values of individual embedding modules (“Int”, “Rel”, and “Ann”) and the combinatorial models based on any two of them (“Int+Rel”, “Int+Ann”, and “Rel+Ann”) for the four types of interactions. The dotted lines indicate the AUC values of the complete DeepERA model.

The radar plot in Fig. 2a showed that the AUC values of all studied approaches for Type 4 interactions were the highest (∼0.88 to ∼0.92), followed by those of Type 3 interactions (∼0.86 to ∼0.9), Type 2 interactions (∼0.7 to ∼0.83), and Type 1 interactions (∼0.6 to ∼0.78), which supports the intuition that the interactions are more difficult to predict when the proteins and/or drugs are unseen in the training set. Besides, the lower AUC values of the cases with unseen proteins (∼0.70 to ∼0.83) compared with the cases with unseen drugs (∼0.76 to ∼0.9) partially reflects the fact that the sequences of proteins are more intricate than the SMILES strings of drugs and therefore more difficult to generalize to new input.

To further investigate the effect of applying different intrinsic embedding modules in DeepERA, we constructed three DeepERA models by, in turn, taking three state-of-the-art deep learning methods, i.e., DeepCPI, DeepDTA, DeepDTI, as their intrinsic embedding component (The model based on DeepCPI is set as the default DeepERA model throughout other analyses in the paper). Replacing the intrinsic embedding model of DeepERA brought little difference to the predictions (Fig. S1) which implied the general dominance of DeepERA over the deep learning methods based on intrinsic features only comes from relational embedding and annotation embedding, especially for the interactions involving unseen proteins (Type 1 and 2 interactions).

### Contributions of the three embedding components in DeepERA

As the DeepERA model is a composite of three components (intrinsic embedding, relational embedding, and annotation embedding), assessing the role of each component in isolation can improve interpretability as to the particular strengths contributing to DeepERA performance. We, therefore, studied the performance of the three individual components for different types of interactions. The results in Fig. 2b showed that the intrinsic embedding had the highest AUC value (0.8663) for the Type 3 interactions and the lowest AUC value (0.706) for the Type 2 interactions among the three components, which suggest that the intrinsic embedding properly captured the attributes of small molecules based on the SMILES strings but was not suitable to resolve the complicated protein solely based on 1D sequences.

On the contrary, the relational embedding showed the highest AUC value (0.8399) and had the largest number of unique true positive predictions (Fig. 2b). As shown in the Venn diagram (Fig. 2c), Type 2 interactions identified 188 unique true interactions compared with 44 identified by intrinsic embedding and 15 identified by annotation embedding. These comparisons indicated that it is possible to make an adequate inference of whether a drug and a protein interact with each other based on the interactions on the local network even though their sequence and annotations were not provided to the model. We also observed this trend in Fig. 2d, in which the performances of combination models including relational embedding (“Rel+Ann”: 0.8124, “Int+Rel”: 0.8219, “Rel”: 0.8399) were higher than the models without it (“Ann”: 0.745, “Int+Ann”: 0.752, “Int”: 0.706) for the Type 2 interactions. Besides, we summarized the unique interactions for each module of Fig. 2c in Table S1 and found that relational embedding tends to identify a comprehensive list of drugs interacting with the unseen proteins.

Annotation embedding showed similar (or slightly better given the proteins are unseen) performance with intrinsic embedding in Fig. 2b. But the combinatorial model of these two components (“Int+Ann”) and the complete DeepERA model achieved higher AUC values (“Int+Ann”: 0.7391, DeepERA: 0.7852) compared with any individual of them (“Int”: 0.679, “Ann”: 0.7078) for the Type 1 interactions (the first radar plot in Fig. 2d), which demonstrated the necessity of annotation embedding in our architecture.

### Our unique protein and drug networks are vital for the relational embedding

The intuition behind the studies utilizing the relationship between drugs and proteins to predict drug-protein interactions is the so-called “guilt-by-association” assumption. First described by Wang and his colleagues (Wang et al., 2013), the assumption suggests that highly similar drugs are more likely to target the same protein and has been extended by Li and his colleagues (Li et al., 2016) to show that a drug targeting the majority of a protein’s neighbors is likely to target that protein and vice versa. Together these imply that adjacent proteins/drugs are likely to bind with the same drug(s)/proteins(s) making their relational networks highly relevant to the drug-target identification task.

To facilitate the relational embedding, we carefully designed unique protein network and drug network. We constructed our protein network as an undirected graph by connecting all pairs of proteins sharing at least one domain in common. We used the drug-drug interactions (DDIs) described and categorized in DrugBank (Wishart et al., 2018) as the relationship in our drug network. Among dozens of categories of DDIs, we selected those that are described as “affect the risk or severity of side effects, therapeutic efficacy, or serum concentration of drugs” and that are not described as “interfere with liberation, absorption, distribution, metabolism, or excretion processes”. (See more details in Method section). To evaluate whether the networks we constructed were compatible with the “guilt-by-association” assumption the relational features were based in, we calculated the proportion of connected drugs that targeted at least one common protein or vice versa. We then compared the proportions from our networks with those from fully randomized networks (drug or protein nodes are randomly connected) (Fig 3a,c) and found that our drug and protein networks were enriched for pairs sharing a common protein or drug interaction. Specifically, the odds ratios are 7.1 (*p*_*val*_ < 1*e*^-200^) and 1.4 (*p*_*val*_ < 1*e*^-140^) for protein networks and drug networks, respectively, which demonstrated our unique networks support the assumption of “guilt by associations”. We also compared the unique network construction against other available networks and the results showed that our protein network and drug network have higher proportions of connected nodes sharing reported drug-target interactions (Fig. 3b,d).

**Fig. 3:**
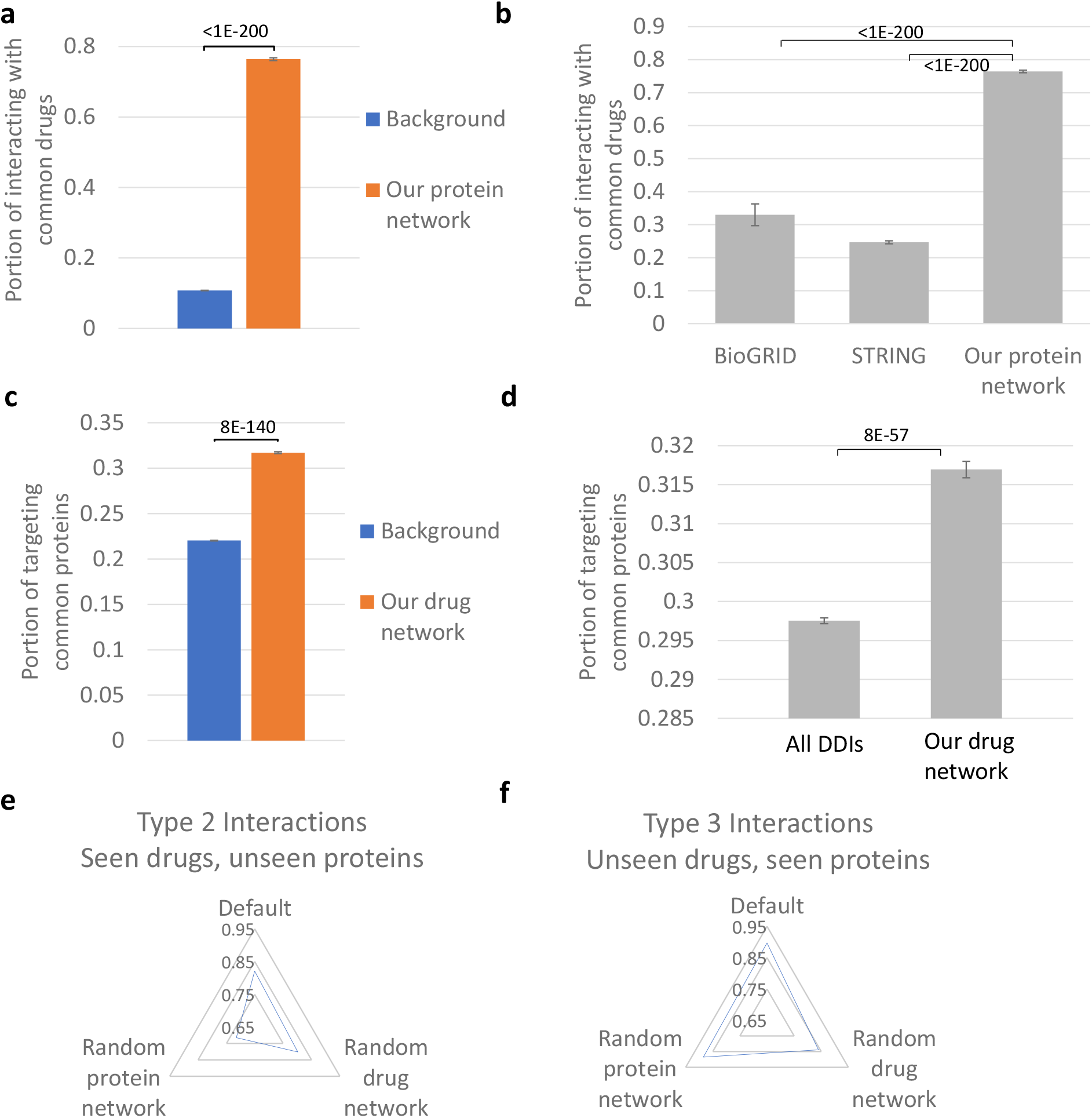
Study the contributions of our unique networks on the predictions. **a** Comparison of portions of connected proteins interacting with at least one common drug in our protein network and the randomized network (background). **b** Comparison of portions of connected proteins interacting with at least one common drug in our protein network and other popular protein networks. **c** Comparison of portions of connected drugs targeting at least one common protein in our drug network and the randomized network (background). **d** Comparison of portions of connected drugs targeting at least one common protein in our drug network and the network based on all DDIs in DrugBank. Comparisons of AUC values between the model using our networks and the models using randomized protein network/drug network for Type 2 interactions (seen drugs and unseen proteins) and Type 3 interactions (unseen drugs and seen proteins) are shown in **e** and **f**, respectively.

We further investigated the importance of our unique protein network and drug network on the performance of DeepERA by replacing the networks with the corresponding random or identity (no connection) networks with the same number of connections (edges). The radar plots showed that the AUC value of DeepERA decreased by 0.013-0.106 (1.42%-12.91%) when the networks are randomized (Fig. 3e,f), especially in the situation that the testing proteins are unseen and the protein network is randomized (Fig. 3e), which brought a decrease of 0.106 (12.91%) for the AUC value of DeepERA. These results implied that our unique networks are good sources of relational information and can theoretically improve the prediction of drug-protein interactions in the DeepERA architecture.

### Identifying drugs for the whole human proteomes and the human proteins interacting with SARS-CoV-2 proteins

We applied the well-trained DeepERA to drug repurposing for 12,920 human proteins. We used 0.5 (the standard threshold) as the first decision cutoff to generate the list of predicted interactions with medium confidence and 0.69, which was an “elbow” point larger than the standard threshold in the curve of distribution of prediction numbers in Fig. S2, as the second decision cutoff to generate the list of high confidence predictions. As a result, we made 28,990,235 predictions for all possible pairs between the studied drugs and 12,954 human proteins and identified 45,603 novel drug-protein interactions with high-confidence (Table S5). Among all 12,954 proteins, we focused on the targets and potential targets of FDA-approved drugs (in total 1,598 proteins listed in The Human Protein Atlas (Uhlén et al., 2015)) and visualized the predicted interactions in a graph (Fig. 5). The graph shows most of the targets/potential targets are related to the diseases about Urinary system, Nervous system, Congenital disorders of metabolism, and Congenital malformations.

We also applied the model to drug repurposing for COVID-19 by predicting the new drugs targeting human proteins known to interact with SARS-CoV-2 viral proteins. There were 332 high-confidence protein interactions between SARS-CoV-2 proteins (bait) and human proteins (prey) identified by the AP-MS analysis (Gordon et al., 2020). To make full use of DeepERA, we excluded the candidate prey proteins without relational information in our work or that had already been seen during the training of the DeepERA model (19 proteins). Finally, we identified 34 proteins for which the DeepERA predictions identified one or more potential interactions within the existing drugs set. Among which, we obtained 344 predicted high-confidence interactions between the studied drugs and 34 human proteins that interacted with SARS-CoV-2 proteins (Table S2).

To provide further support our predictions, we performed computational docking for a limited number of cases with medium or high prediction scores with Autodock Vina (Oleg Arthur J., 2010). After excluding unavailable proteins and drugs (e.g., have no PDB structure or cannot be processed by Autodock Vina), we obtained 846 docking results for 14 prey proteins, which have been appended to the prediction lists. Autodock Vina, by utilizing an empirical scoring function inspired by the X-score function, generated a negative computational score (affinity) along with the docking result and this negative score represents the free energy of the binding pose between the drug and the target (the lower the better). However, it is well known that there are no general guidelines to determine the confidence of interactions based on docking scores. Besides, some proteins/drugs might be easier to get a highly negative score (i.e., low affinity values) in the docking, meaning the docking scores of different proteins/drugs cannot be directly compared. To deal with these problems, we applied a two-way test (see Methods section) to “normalize” the docking scores and found 12 predicted interactions with significant scores (Table S2).

Fig. 4a summarized the computational docking results obtained by using AutoDock Vina (DeLano, 2002) for 11 prey proteins with all drugs in this study. The docking results with non-negative affinities and failed docking results are neglected. To find the most interesting cases, we designed a way to test the significance of the docking results (see Methods in Supplementary Notes) and identified 12 drug-target pairs with significantly high affinities (Fig. 4a). From the 11 prey proteins, we selected TLE1 as the target for further study. TLE1 can bind to many transcription factors and repress their ability to bind DNA. It is also an innate immune signaling protein in the NF-κB pathway which is targeted by the viral protein NSP13 of SARS-CoV-2 (Gordon et al., 2020). Importantly TLE1 had not yet been recognized as a drug target prior to this study. For this potential SARS-CoV-2 target protein, we identified 7 drugs interacting with it. Venetoclax and Foretinib are the drugs with the best docking results with TLE1 (PDB entry: 2CE8) and their docking affinity values are significantly lower than the corresponding random combinations (Fig. 4b-e). The receptor-ligand complexes of TLE1-Venetoclax> and TLE1-Foretinib are visualized by PyMol (Fig. 4f,g).

**Fig. 4:**
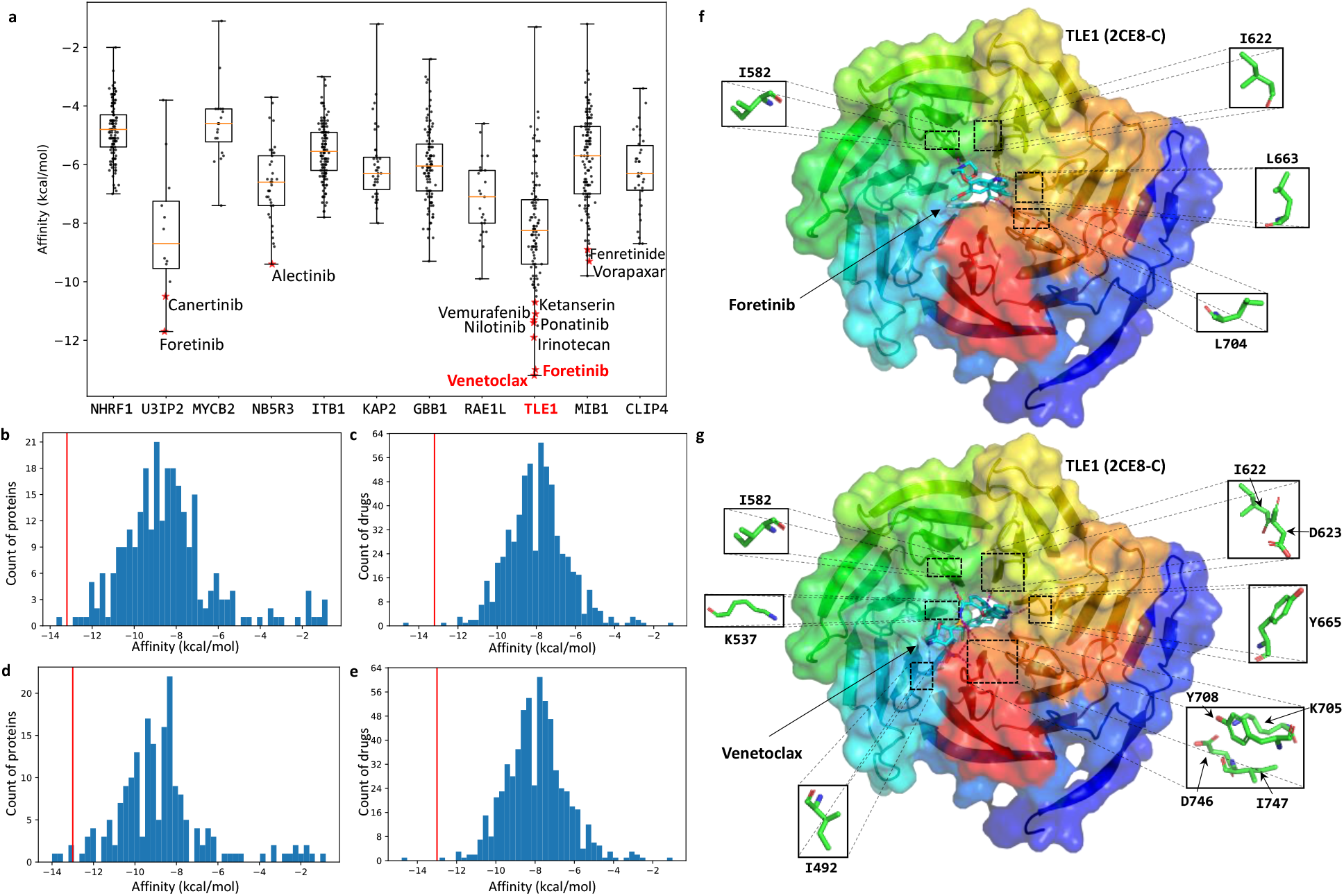
Normalized docking result of predicted drug-protein interactions. **a** Docking affinities of predicted interactions between 11 human prey proteins of SARS-CoV-2 and the selected drugs. The interactions with significant normalized docking affinities are emphasized by red asterisk mark (*). Interactions of <TLE1, Venetoclax> and <TLE1, Foretinib> are selected for further study. **b** Distribution of docking affinity between Venetoclax and random proteins. **c** Distribution of docking affinity between TLE1 and random drugs. Affinity of <TLE1, Venetoclax> is indicated by red vertical lines in both **b** and **c. d** Distribution of docking affinity between Foretinib and random proteins. **e** Distribution of docking affinity between TLE1 and random drugs. Affinity of <TLE1, Foretinib> is indicated by red vertical lines in both **d** and **e**. The docked receptor-ligand complexes of <TLE1, Venetoclax> and <TLE1, Foretinib> (PDB entry of TLE1: 2CE8) are visualized by open-source PyMol in **f** and **g**, respectively. Only chain-C of 2CE8, to which Venetoclax and Foretinib bound, is included in the visualization.

**Fig. 5:**
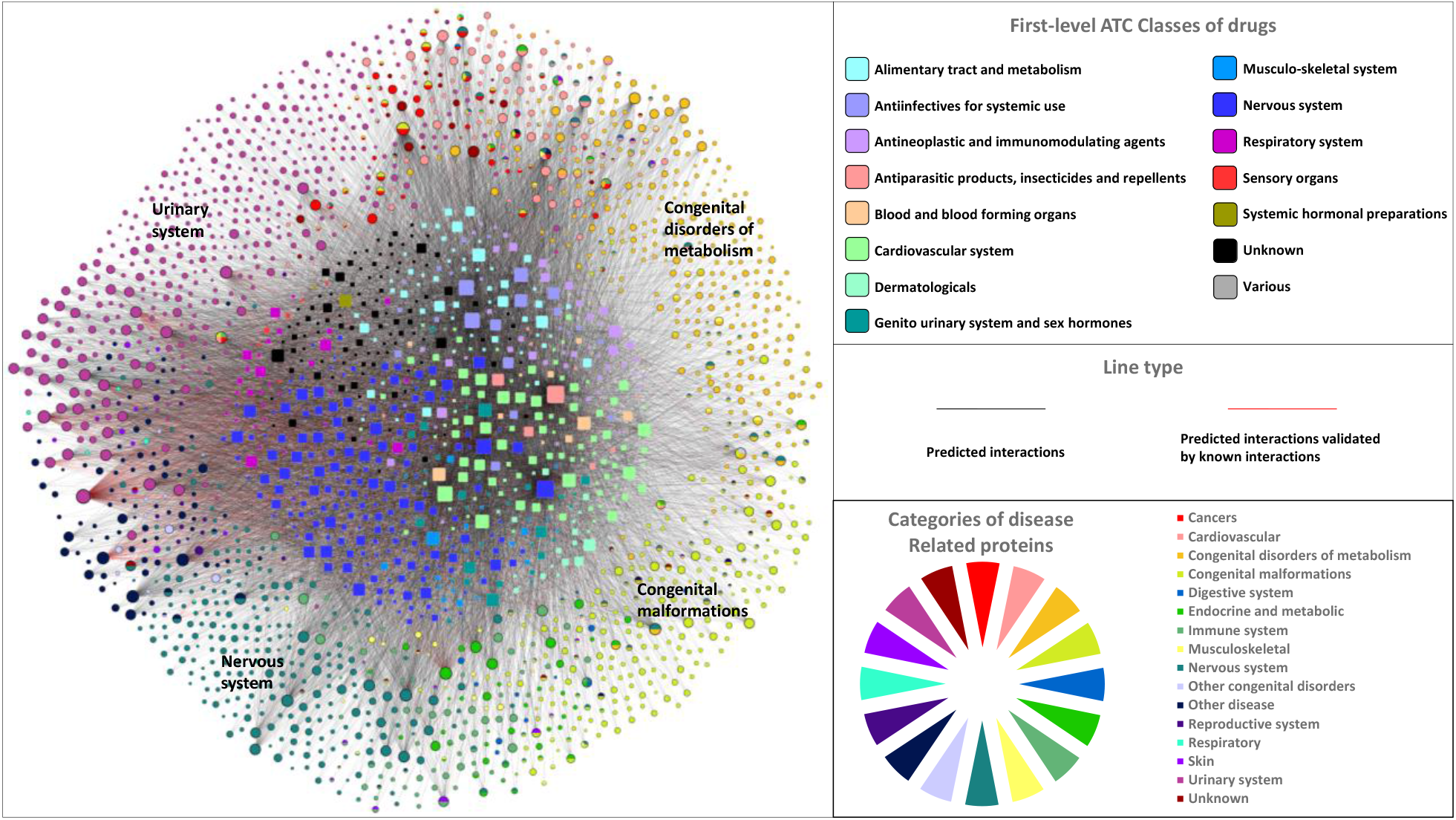
A drug-protein bipartite graph predicted by DeepERA. A drug node (square) and a protein node (circle) are connected to each other by an edge if the protein is predicted to interact with the drug by DeepERA. Edges are colored as red if they are annotated to have known experimental interactions in DrugBank. Drug nodes are colored based on their first-level anatomical therapeutic chemical (ATC) classification. Protein nodes are colored based on the categories of related diseases in The Human Protein Atlas and pie charts are used for the proteins with multiple category labels. In this graph, only the proteins considered as the targets of FDA-approved drugs or potential targets identified by The Human Protein Atlas are included. The graph was prepared by Cytoscape (http://www.cytoscape.org/).

## Discussion

In this paper, we proposed a deep learning-based embedding architecture, named DeepERA, that learned comprehensive features of drugs and proteins from their intrinsic attributes (SMILES strings/sequences), relationship (drug-drug/protein-protein networks), and annotation profiles (disease/side effects and annotations) and provided comprehensive identification of drug-protein interactions. This is the first work that combined the advantages of deep learning and heterogeneous data to facilitate the prediction of drug-protein interactions.

Our study showed that DeepERA has better performance than other existing deep learning embedding approaches. To explore the reason for this, we studied the performance of three individual embedding components applied in DeepERA and found that each embedding module has its preference, which enabled the improvement of the overall predictive performance of DeepERA in which all three modules are combined. Considering the high substitutability of the embedding modules of our architecture, any future replacement with one or more advanced embedding module(s) will potentially further improve DeepERA’s predictive performance.

We also found that our unique relational networks played important roles to the performance of the relational embedding model. With a trained DeepERA model, we generated a list of potential drugs for the human proteins that were identified as interacting with SARS-CoV-2 viral proteins and provided computational docking with the significance of results determined by statistical tests to support the predictions.

One recent work (Lim et al., 2019) investigated the compatibility of deep learning models with drug-target prediction based on the 3D structure of proteins and drug molecules. It demonstrated that deep learning models are also capable of identifying drug-protein interactions based on the 3D structure, which can also contribute to the predictions. Due to the limited numbers of available 3D data (only 72 proteins and 25 proteins in the training and testing set, respectively), though, we did not add it into the current version of DeepERA. But the recent earthquake in protein structure prediction (protein folding problem), the development of AlphaFold2 (Jumper et al., 2021), which has been considered as a solution to the problem, brings us the hope of having a large amount of protein 3D structures in a near future. At that time, the DeepERA model can be easily extended to accommodate 3D structure information for more comprehensive and accurate predictions of drug-protein interactions.

---

**Fig. S1:**
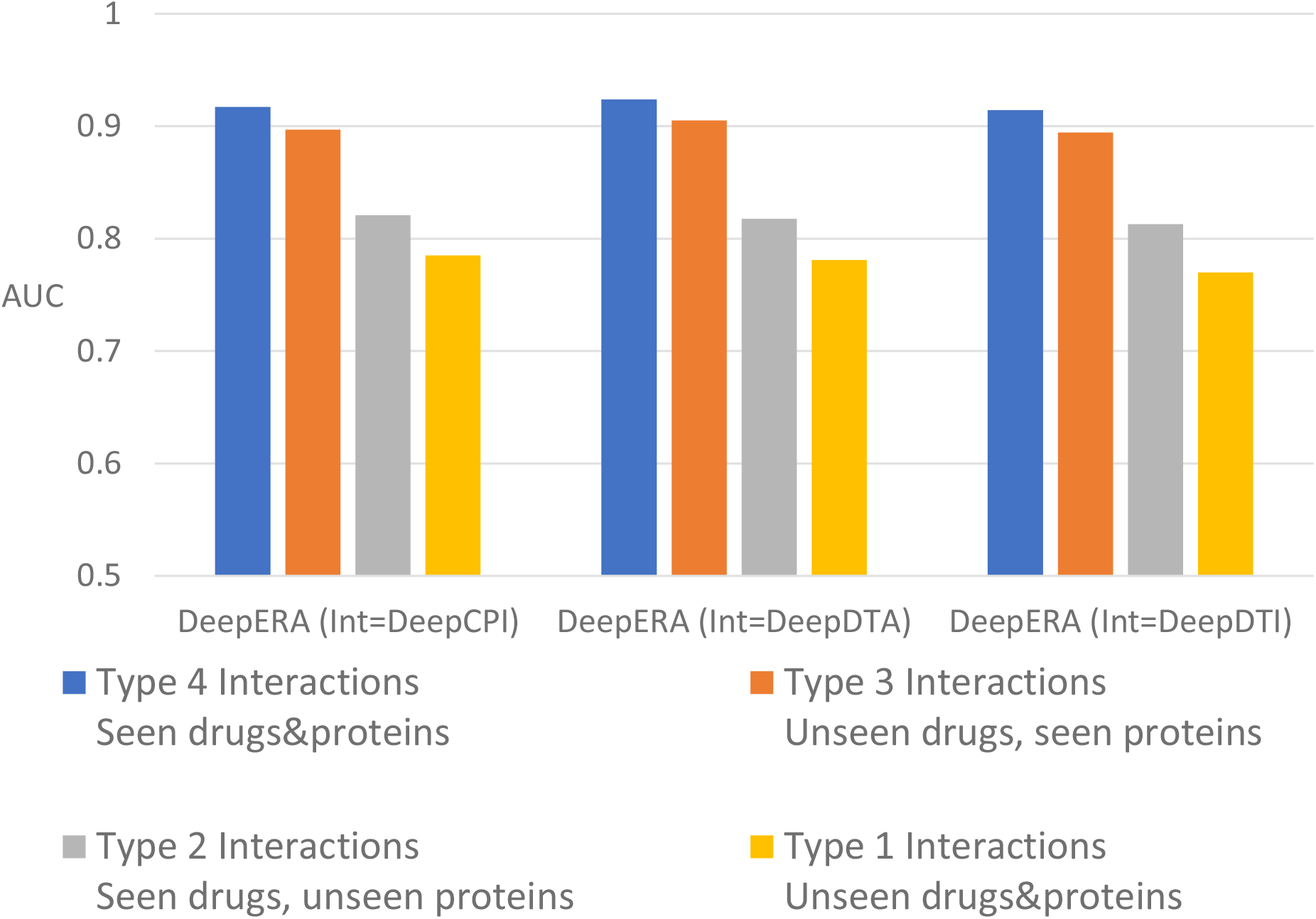
Performance of DeepERA with different intrinsic embedding models (Int = DeepCPI, DeepDTA, DeepDTI).

**Fig. S2:**
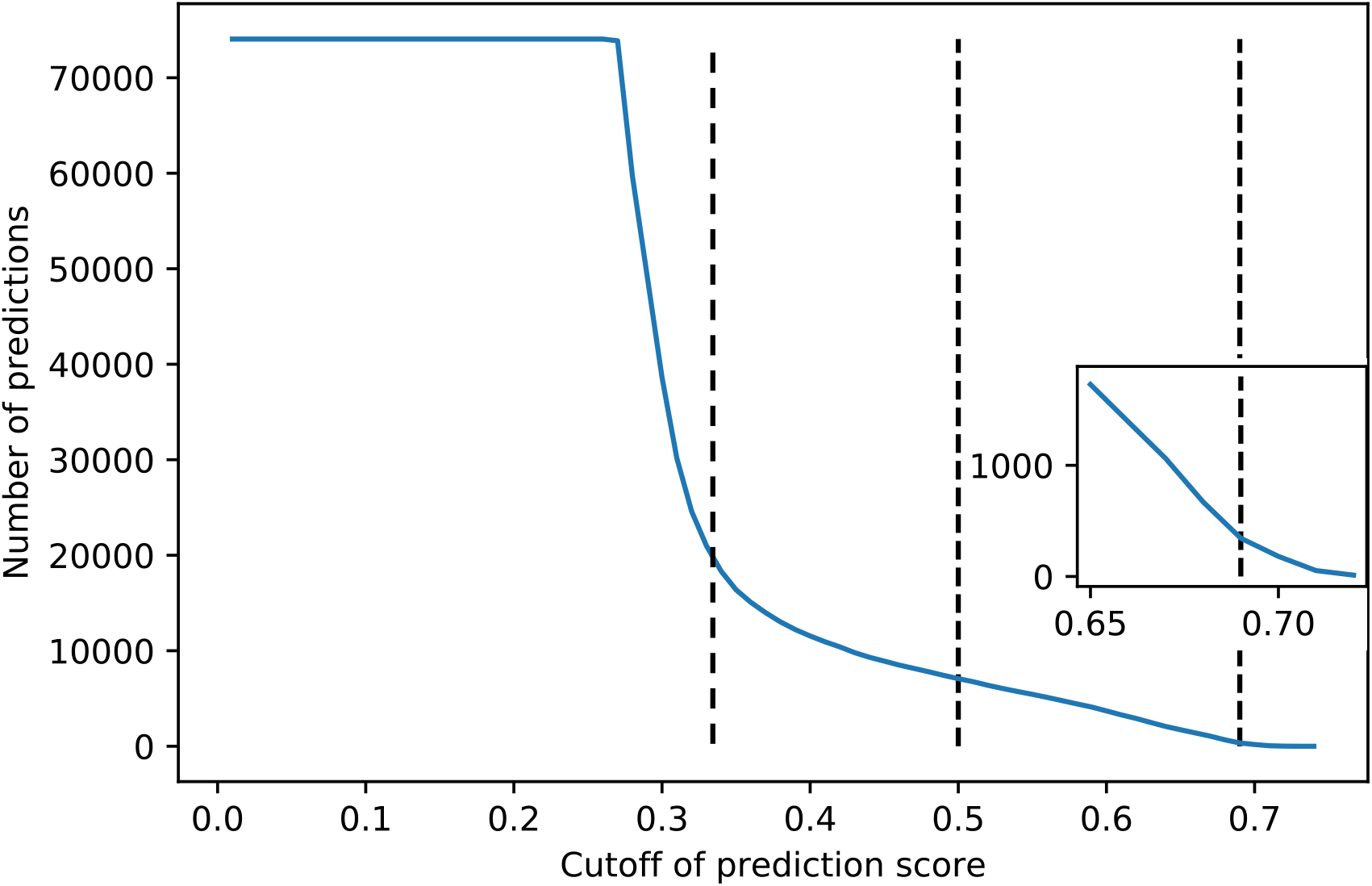
Distribution of number of predictions along with the score cutoff. The low-confidence, medium-confidence, and high-confidence cutoffs at 0.33 (the first elbow point), 0.5 (default cutoff in classification), and 0.69 (the second elbow point) are indicated by dashed lines. The elbow points are the points on the curve with locally maximal absolute second derivative and using elbow points as cutoff is a common heuristic in optimization.

**Fig. S3:**
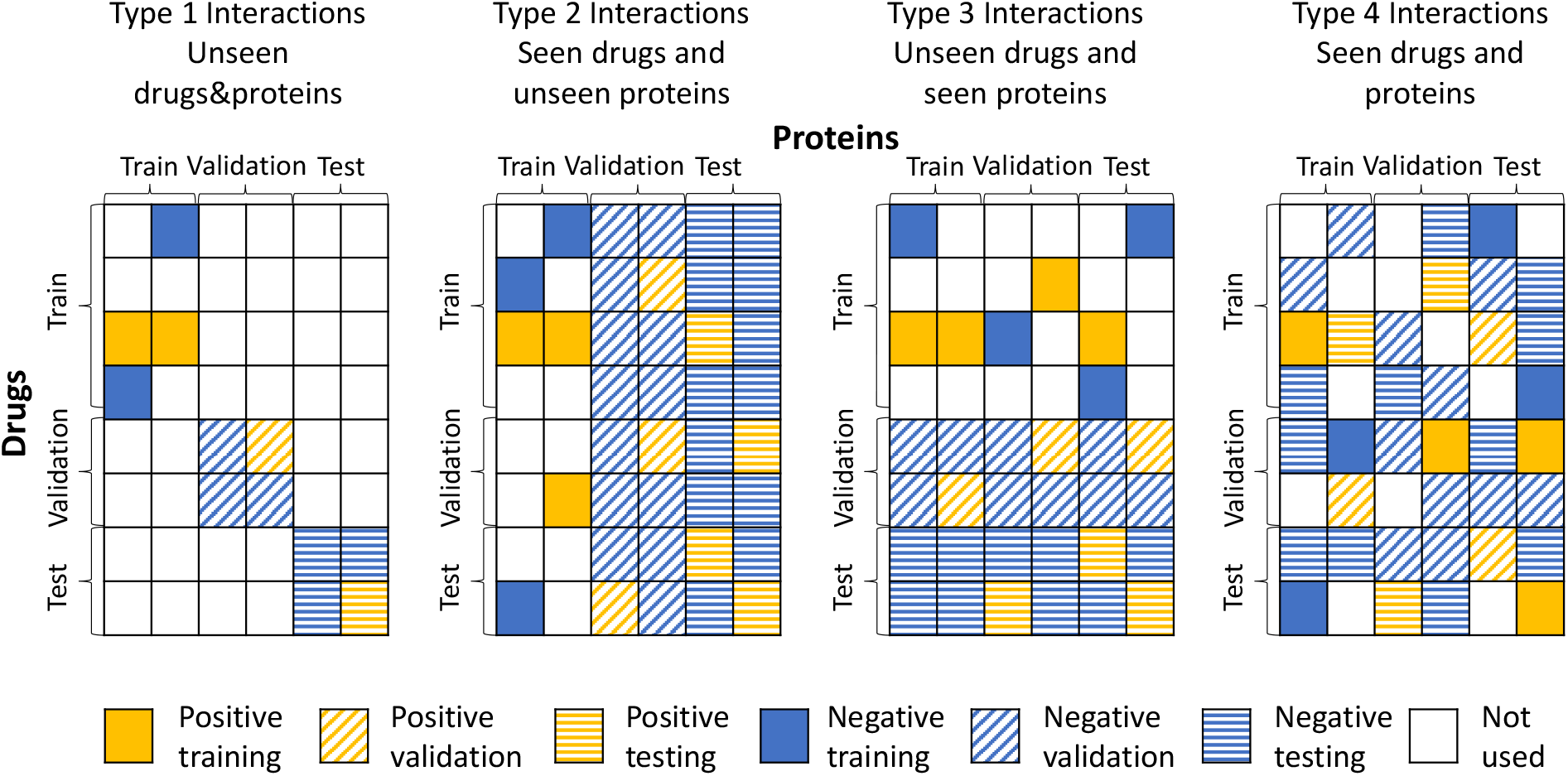
Separation of training, validation, and testing samples for the four types of interactions.

**Fig. S4:**
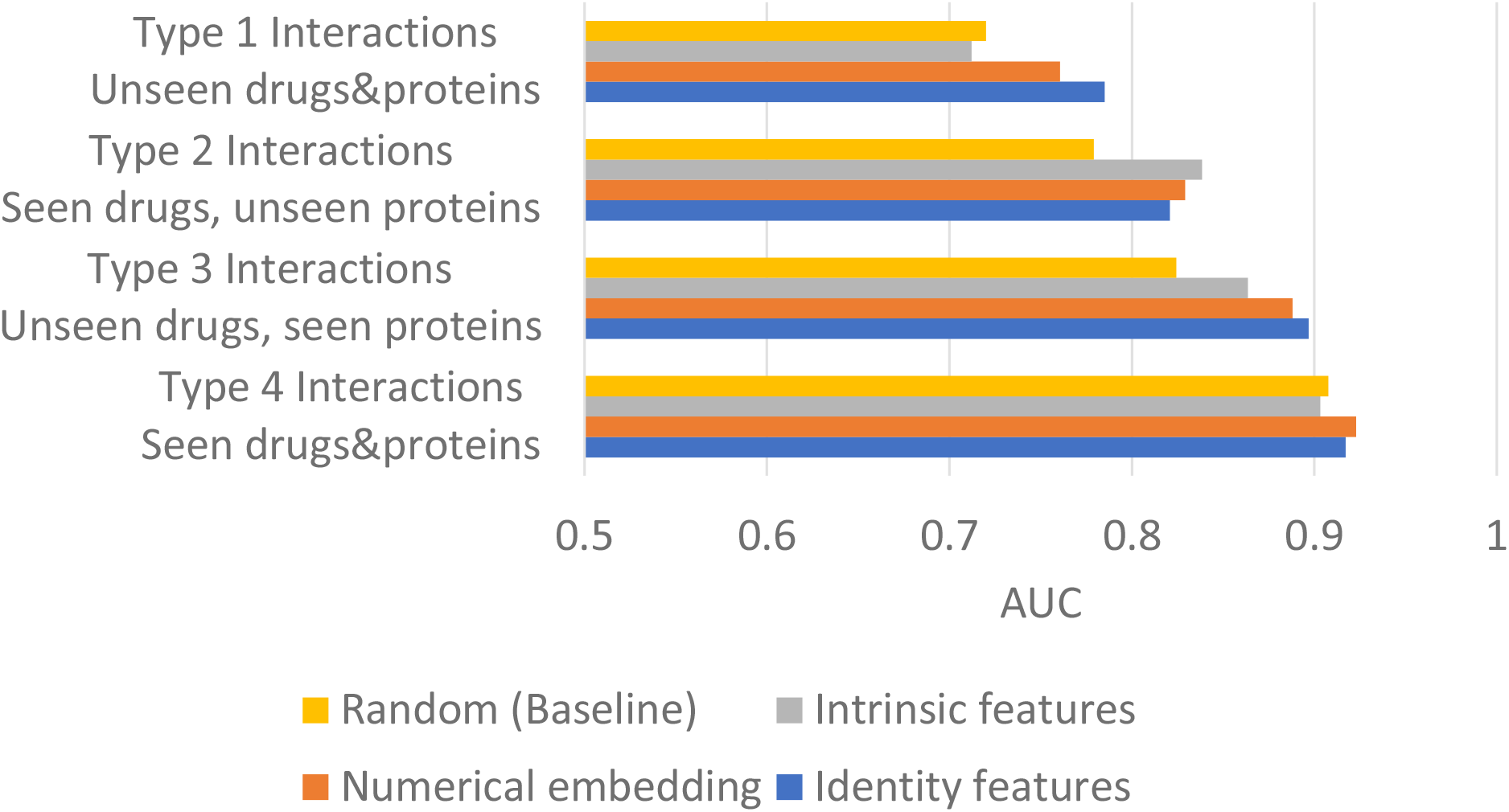
The comparison of using different initialization features for drugs and proteins in the graph neural network of relational embedding. The result of using randomly initialized features is taken as the baseline for the comparison.

**Fig. S5:**
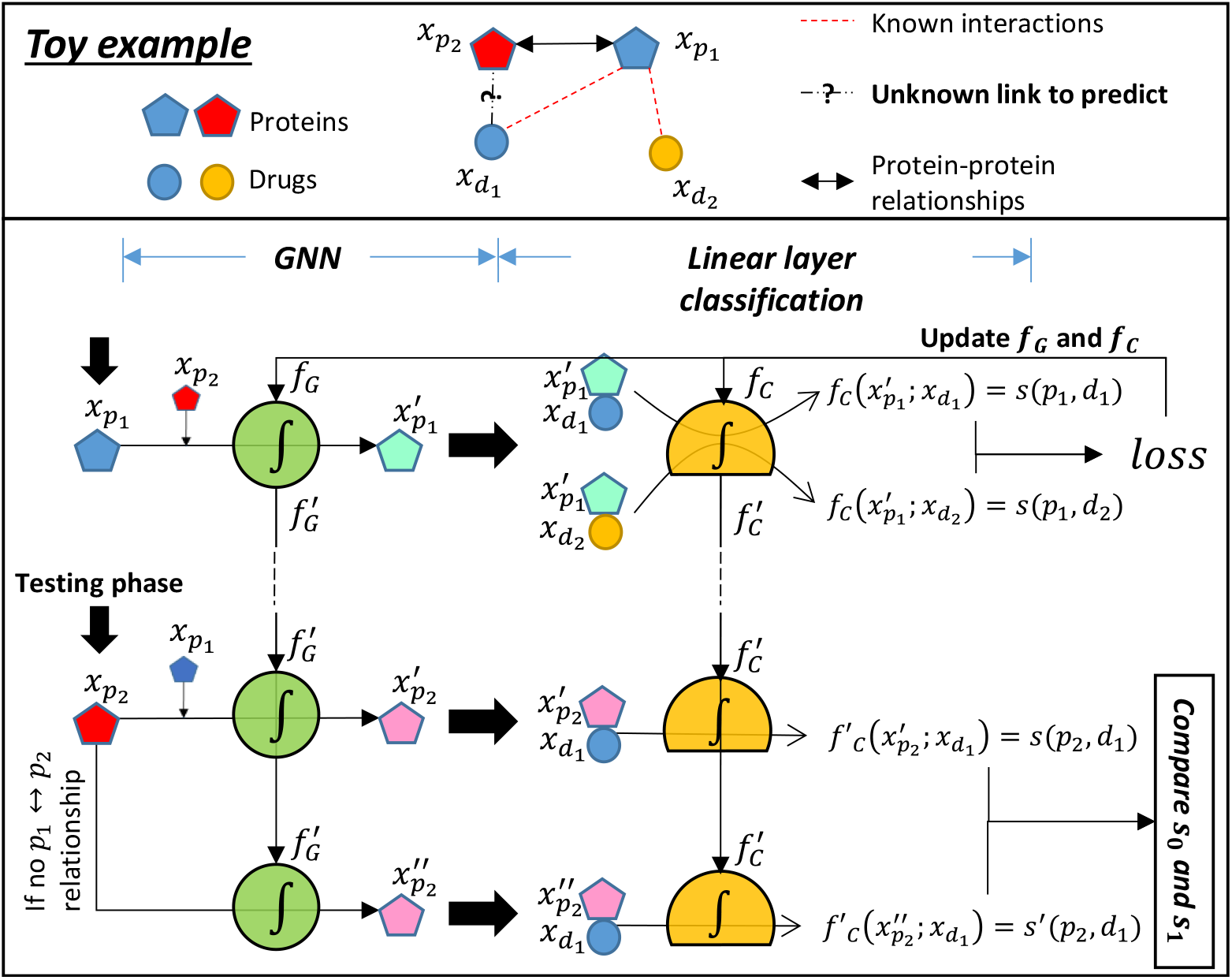
A toy example of explaining the mechanism of relational embedding based on graph neural network.

